# Bioinformatics Study on Structural Proteins of Severe Acute Respiratory Syndrome Coronavirus 2 (SARS-COV-2) For Better Understanding the Vaccine Development

**DOI:** 10.1101/2020.04.21.053199

**Authors:** Sumaira Gulzar, Saqib Husssain

## Abstract

Novel coronavirus 2019 (2019-nCoV), also known as SARS-CoV-2), leads high morbidity and mortality in global epidemics. Four structural proteins (surface glycoprotein (QIQ22760.1), envelop glycoprotein (QIQ22762.1), nucleocapsid phosphoprotein (QIQ22768.1) and membrane glycoprotein (QIQ22763.1)) of SARS-CoV-2 are extracted from the NCBI database and further analyzed with ExPASy ProtParam tool. Lucien is the highest in envelope, surface and membrane glycoprotein that is an optimal environment for rapid virus fixation on host cell's surface to the receptor molecule. Transmembrane region prediction was performed by SOSUI server. For all structural proteins, except nucleocapsid Phosphoprotein, the trans-membrane prediction indicates that the virus can enter the host easily. Domain analysis was done by SMART tool. Domain information helps in the function of the viral protein. Lastly, the 3D structure prediction was carried out by Swiss Model and the result validation was achieved by PROCHECK. Such models are the starting point of the community for structural drug and vaccine designs as well as virtual computational screening.

## I. INTRODUCTION

The latest 2019–nCoV, now officially known as Severe Acute Respiratory Syndrome Coronavirus 2 (SARS-CoV-2), was established as the responsible pandemic. Infections of (SARS-CoV-2) have been reported in more than 200 countries including Pakistan [1]. The first genome sequence of SARS-CoV-2 to be published from Pakistan and is now available on NCBI https://www.ncbi.nlm.nih.gov/nuccore/MT240479, GISAID and NEXTRAIN.

Coronaviruses, a genus belonging to the Coronaviridae family, contain the largest genome (26.4 kb to 31.7 kb) of RNA viruses with a diameter of 120–160 nm, consisting of a single-stranded positive-sense RNA molecule **[2]** The broad genome has given this family of viruses extra plasticity in their adaptation and alteration of genes. The G + C content of coronavirus genomes ranges from 32% TO 43% The genome consists of seven genes arranged in sequence [5′-replicase ORF1ab, spike (S), envelope (E), membrane (M), nucleocapside (N)-3′] with small untranslated regions in both termini and additional ORFs in each subgroup of coronavirus. **[3, 4]**

5 ‘Non-structural protein coding regions consisting of two-thirds genome replicase genes 1 and structural and non-essential gene. 2-7 regions consisting of structural and non-essential protein coding regions **[5]**. The genes are translated from genomic mRNA 2-7. Subgenomic RNAs encode the main Surface Proteins (S), Envelope Protein (E), Membranes Protein (M), and Nucleocapside Protein **[6]**.

Surface protein of SARS CoV-2 join ACE2 (angiotensin-converting enzyme2) and to infect cells. After this initial process, Surface Protein must be produced with an enzyme known as protease to complete entry into the cell. SARS CoV-2 uses TMPRSS 2 in the same way as SARS-CoV is used to complete this process **[8]**

Surface Glycoproteins are outside of the virion and give the typical shape to the virion. The S proteins form homotrimers that allow sun-like morphologies to be developed that give the name Coronaviruses via the C-terminal transmembrane regions, S proteins bind to the virion membrane and interact with M proteins. Virions can be attached through the N-terminus of the S proteins to different surface receptors in the host cell's plasma membrane. The S protein is the receptor-binder and viral input in host cells and is therefore a major therapeutic objective **[9]**.

Membrane Glycoproteins proteins inside the Golgi system are glycosylated. The modification of the M protein is essential for the virion to attach into the cell and make antigenic protein **[10]**.

The protein M plays a key role in the cell's regeneration of virions. N protein forms a complex by binding to genomic RNA and M protein activates the development of interacting virions in this intermediate endoplasmic reticulum-Golgi interface (ERGIC) compartment with this complex **[11,12].**

**Figure 1.**
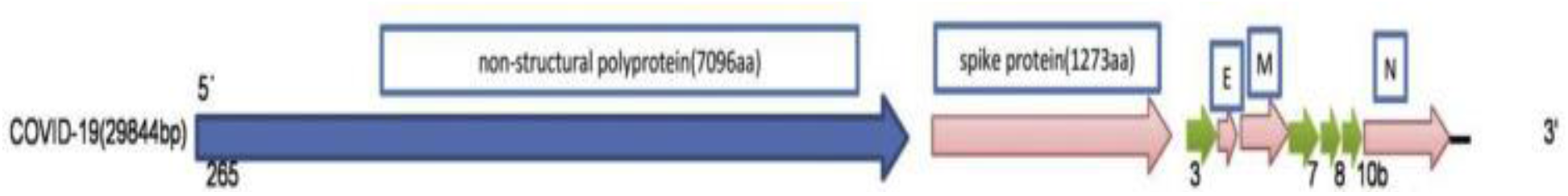
The COVID-19 region of 5′ UTR and 3′ UTR and coding, the number of base pairs shown. [7].

Envelop Glycoproteins are tiny proteins consisting of around 76 to 109 amino acids. Approximately 30 amino acids in the E protein N-terminus allow attachment to the virus membrane **[13]**.

Additionally, coronavirus E proteins play a critical role in virion assembly and morphogenesis within the cell. For one research coronavirus E and M proteins were expressed for conjunction with vectors of mammalian expression to form virus-like structures inside the cell **[14, 15]**.

In another review, the ability of recombinant mouse hepatitis virus (MHV) and SARS viruses to generate E protein expression in the genome to sustain this status has been significantly reduced **[16, 17]**.

N Proteins are helix-capable phosphoproteins with a versatile viral genomic RNA structure. It plays an important role in coronavirus virion assembly, replication and transcription, as the N protein locates both the coronavirus replication / transcription region and the ERGIC region where the virus is collected **[18]**. In this paper, we tried to describe the four structural proteins of SARS-CoV-2 by the use of bioinformatics tools

## II. MATERIALS AND METHODS

Four structural proteins of SARS COV-2 included in this research named as surface glycoprotein (QIQ22760.1), envelop glycoprotein (QIQ22762.1), nucleocapsid phosphoprotein (QIQ22768.1), membrane glycoprotein (QIQ22763.1) available in the NCBI was retrieved. For further study of bioinformatics, the FASTA sequence was selected and used.

### Protein statistics

Using the Protparam method, protein statistics were reviewed. ProtParam method enables the measurement of various physicochemical protein sequence parameters. Parameters included molecular weight, Theoretical Pi (Isoelectric point), Half-life calculation, instability index, aliphatic index and Grand range of hydropathicity (GRAVY)

### Trans-membrane sequence prediction

SOSUI tools from EXPASY repository are used for predicting transmembrane. Results normally return within 1 min. It provides the following production number of TMHs with sequence and protein type

### Functional Site Prediction

One of the main goals of molecular biology is functional assignment to protein. Protein functional sites that are responsible for or perform all of the essential protein functions are more conserved than other regions over the time of evolution. These accessible websites are called domains. SMART Internet server was forecasting domains

### 3D Model predictions and validation

The SARS COV-2 structural protein amino acid sequences are used as targets for homology modeling using the SWISS-MODEL server. Depending on the Global Model Quality Estimation (GMQE) and QMEAN, the top-ranked models are further analyzed and sorted. GMQE is a quality estimate that combines the target – template alignment properties with the template search method. The corresponding GMQE value is given as 0 to 1. The QMEAN Z-score gives a global estimation of the “degree of nativeness” of the structural characteristics observed in the model and is described in Benkert et al. **[19]** QMEAN Z-scores about zero indicate strong compatibility between the structure of the model and similar-size experimental structures. Scores of −4.0 or below are indicative of poor quality models. The overall stereochemical output was checked by PROCHECK including torsional angles to the backbone through the Ramachandran plot **[20]**

## III. RESULTS

### Protein Statistics

Phenyl alanine and serine are absolutely missing in all proteins tested as displayed in chemical parameters (Table 1). Histidine and glutamic acid is absent in envelope protein and cystein lacks in nucleocapsid phosphoprotein. lucien quantity is more in envelop protein, surface and membrane glycoprotein while glycine is more in nucleocapsid phosphoprotein

**Table 1.** Amino Acid Composition the Structural Protein Extracted From SARS-COV-2.

| Nucleocapsid Phosphoprotein | Membrane Glycoprotein | Surface Glycoprotein | Envelope Protein | Proteins |
| --- | --- | --- | --- | --- |
| 37 | 19 | 79 | 4 | Ala (A) |
| 29 | 14 | 42 | 3 | Arg (R) |
| 22 | 11 | 88 | 5 | Asn (N) |
| 24 | 6 | 62 | 1 | Asp (D) |
| 0 | 4 | 40 | 3 | Cys (C) |
| 35 | 4 | 62 | 0 | Gln (Q) |
| 12 | 7 | 48 | 2 | Glu (E) |
| 43 | 14 | 82 | 1 | Gly (G) |
| 4 | 5 | 17 | 0 | His (H) |
| 14 | 20 | 76 | 3 | Ile (I) |
| 27 | 35 | 108 | 14 | Leu (L) |
| 31 | 7 | 61 | 2 | Lys (K) |
| 7 | 4 | 14 | 1 | Met (M) |
| 13 | 11 | 77 | 5 | Phe (F) |
| 28 | 5 | 58 | 2 | Pro (P) |
| 37 | 15 | 99 | 10 | Ser (S) |
| 32 | 13 | 97 | 8 | Thr (T) |
| 5 | 7 | 12 | 4 | Trp (W) |
| 11 | 9 | 54 |  | Tyr |
| 8 | 12 | 97 | 13 | Val (V) |
| 0 | 0 | 0 | 0 | Pyl (O) |
| 0 | 0 | 0 | 0 | Sec (U) |

The physical parameters (Table 2) show that Surface protein contain highest amino acids (1273), negatively charged residues (110) positively charged residues (103), EC (148960) and molecular weight (141178.47). Higher molecular weight of the surface glycoprotein suggests that its tertiary structure may contain strong amino acid side chains. All three Proteins are basic in nature, excluding surface glycoprotein, as the isoelectric point value is more than 9.The instability index value greater than 40 is considered unstable [21]This is derived from the Instability index surface, membrane and envelop glycoprotein is stable and nucleocapsid phosphoprotein is highly unstable the range of aliphatic index from 33.01 −55.09 that suggests a tendency to be sensitive to a wide range of temperatures**[22]** and GRAVY value shows protein's hydropathicity and whether the nature protein side chains are hydrophilic or hydrophobic **[23]**. Leaving the surface Glycoprotein and nucleocapsid phosphoprotein all other protein are hydrophobic in nature

**Table 2.**
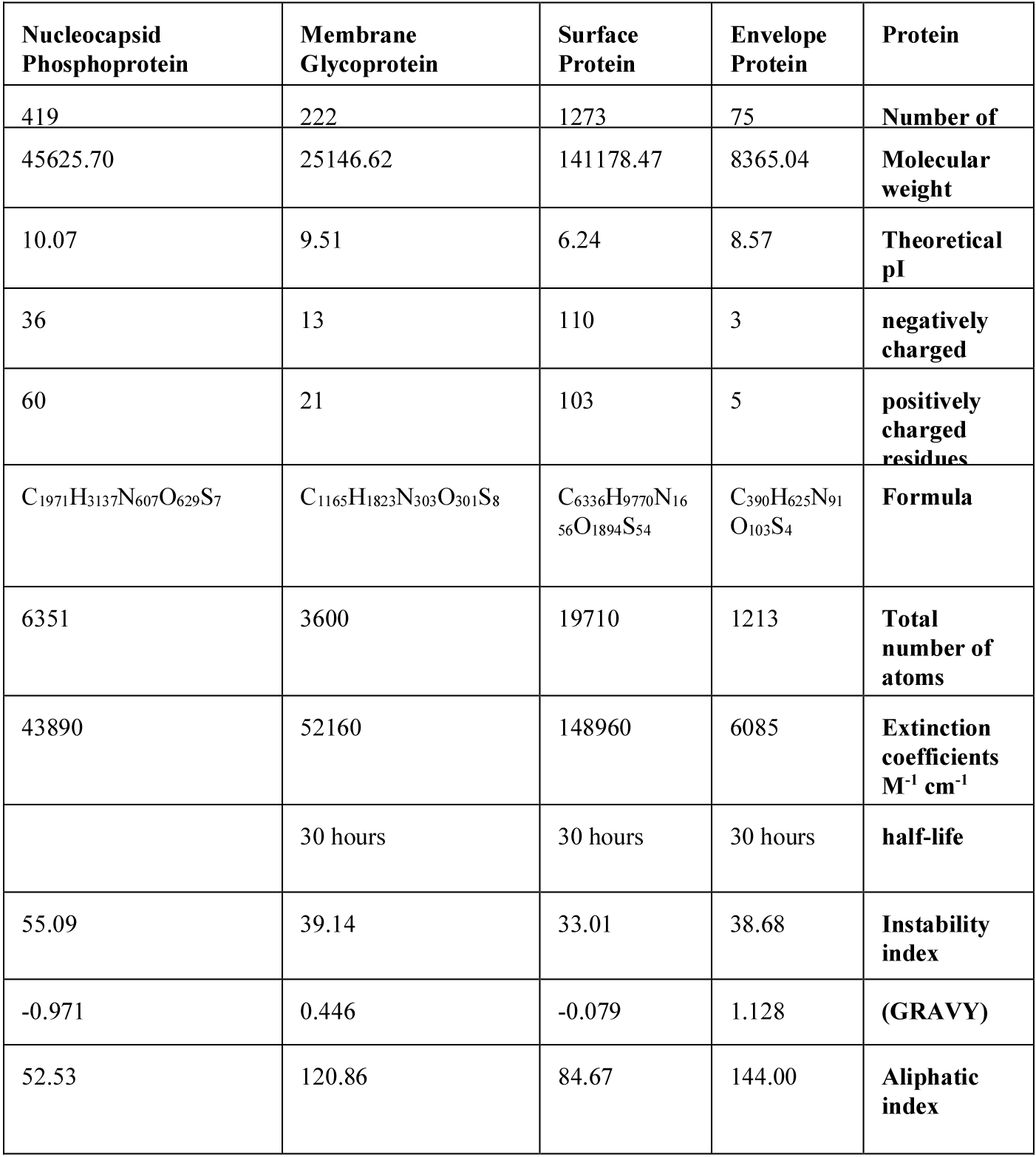
Physical Parameters of the Structural Protein from SARS-COV-2.

### Trans-membrane sequence prediction

Analysis of the transmembrane protein reveals (Table 3) that it is absent in nucleocapsid phosphoprotein. The other three proteins show transmembrane sequence. All three other proteins show different sequence from each other. Membrane glycoprotein have three transmembrane region yet there is a sequence of length 22 and 23 amino acids in all the proteins that show trans-membrane. All the proteins have different C and N terminals

**Table 3.**
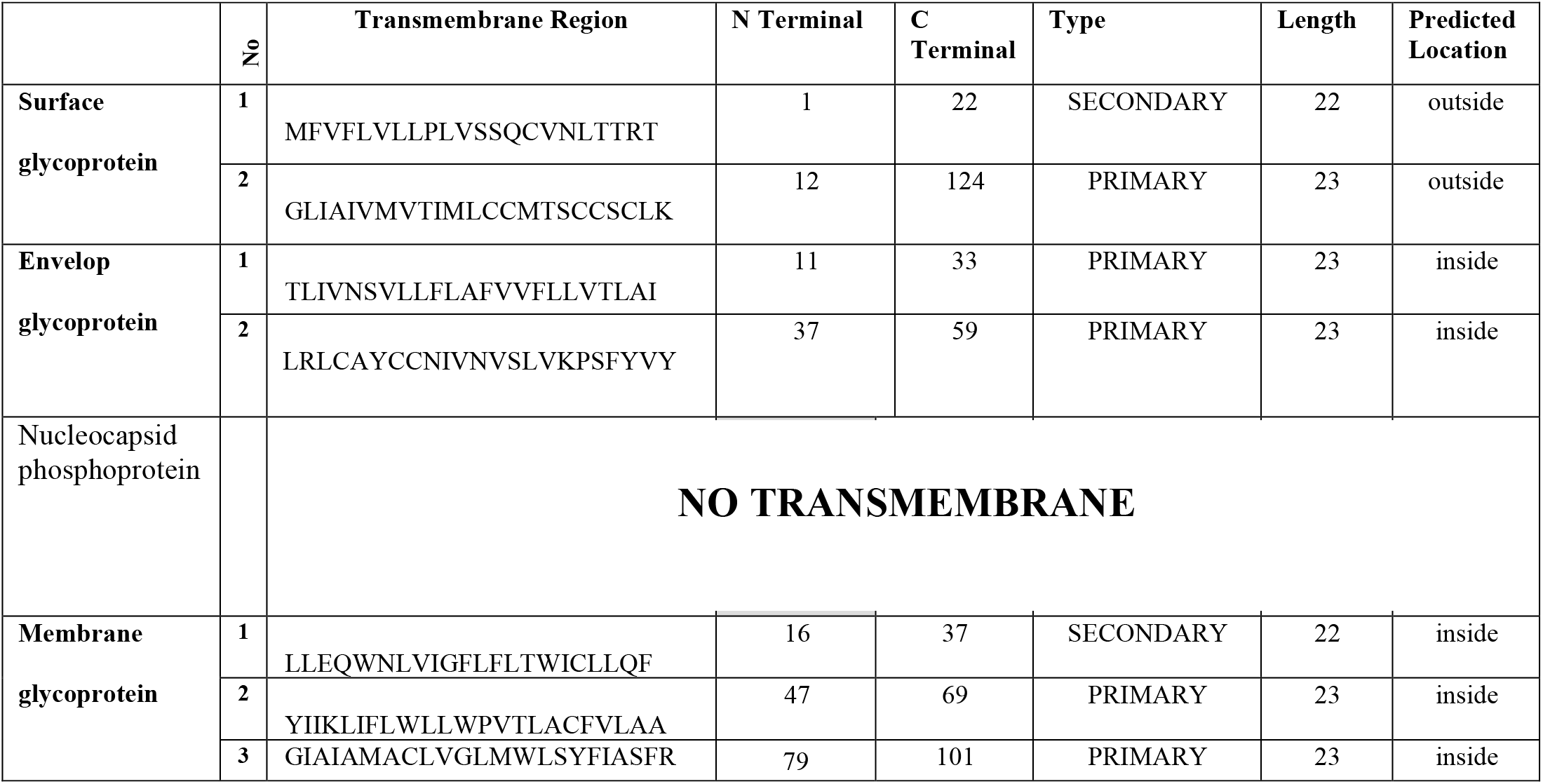
Transmembrane of the Protein Extracted from SARS-COV-2

### Functional Site Prediction

SMART results (Table 4) clearly indicates the Surface Glycoprotein contains two domain. One is Pfam:Spike_rec_bind starts from 330 to 583 residues predicted with the e-value (expected value) second is Pfam:Corona_S2 domain starts from 671 to 1279 residues predicted with the e-value Envelop Glycoprotein contains one domain Pfam:NS3_envE starts from 1 to 74 residues predicted with the e-value 7.7e-8.Membrane Glycoprotein contains one domain Pfam:Corona_Mstarts from 4 to 221 residues predicted with the e-value 1.3e-95.Nucleocapsid Phosphoprotein contains one domain Pfam:Corona_nucleoca starts from 14 to 337 residues predicted with the e-value 1.2e-164 3D Model predictions and validation

**Table 4.** Predicted domains of Structural Proteins of SARS-CoV-2 by SMART.

| Proteins | Domain Name | Start | End | E-value |
| --- | --- | --- | --- | --- |
| Surface Glycoprotein | Low complexity | 2 | 11 | N/A |
|  | Pfam:Spike_rec_bind | 330 | 583 | 6.1e-75 |
|  | Pfam:Corona_S2 | 671 | 1279 | 1.3e-266 |
| Envelop Glycoprotein | Pfam:NS3_envE | 1 | 74 | 7.7e-8 |
| Membrane Glycoprotein | Pfam:Corona_M | 4 | 221 | 1.3e-95 |
| Nucleocapsid Phosphoprotein | Pfam:Corona_nucleoca | 14 | 337 | 1.2e-164 |

### 3D Model predictions and validation

(Table 5 and Figure. 2) shows the evaluation of the 3D model that all four proteins have satisfactory model with QMEAN > 0.Model validity research Shows that the models are at the optimum point, just like in the Ramchandran series.

**Figure 2.**
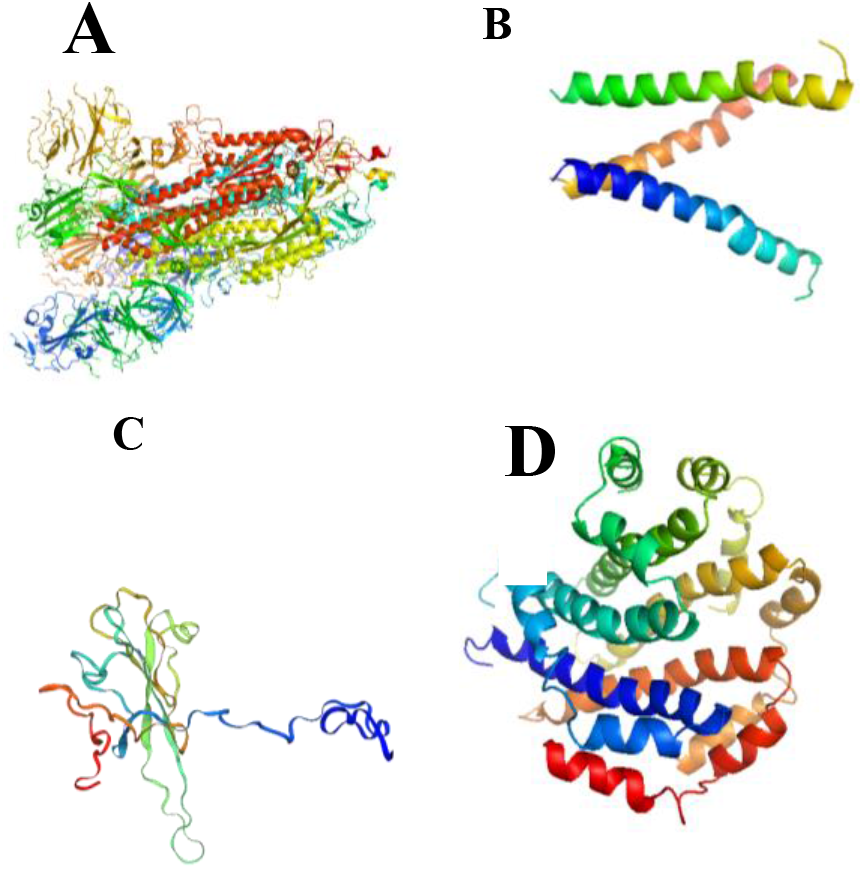
3D Model generated from Swiss Model of SARS COV-2 Structural proteins; Surface glycoprotein (A), Envelop glycoprotein (B), Membrane glycoprotein (C), Nucleocapsid phosphoprotein(D)

(Figure 3).

**Figure 3.**
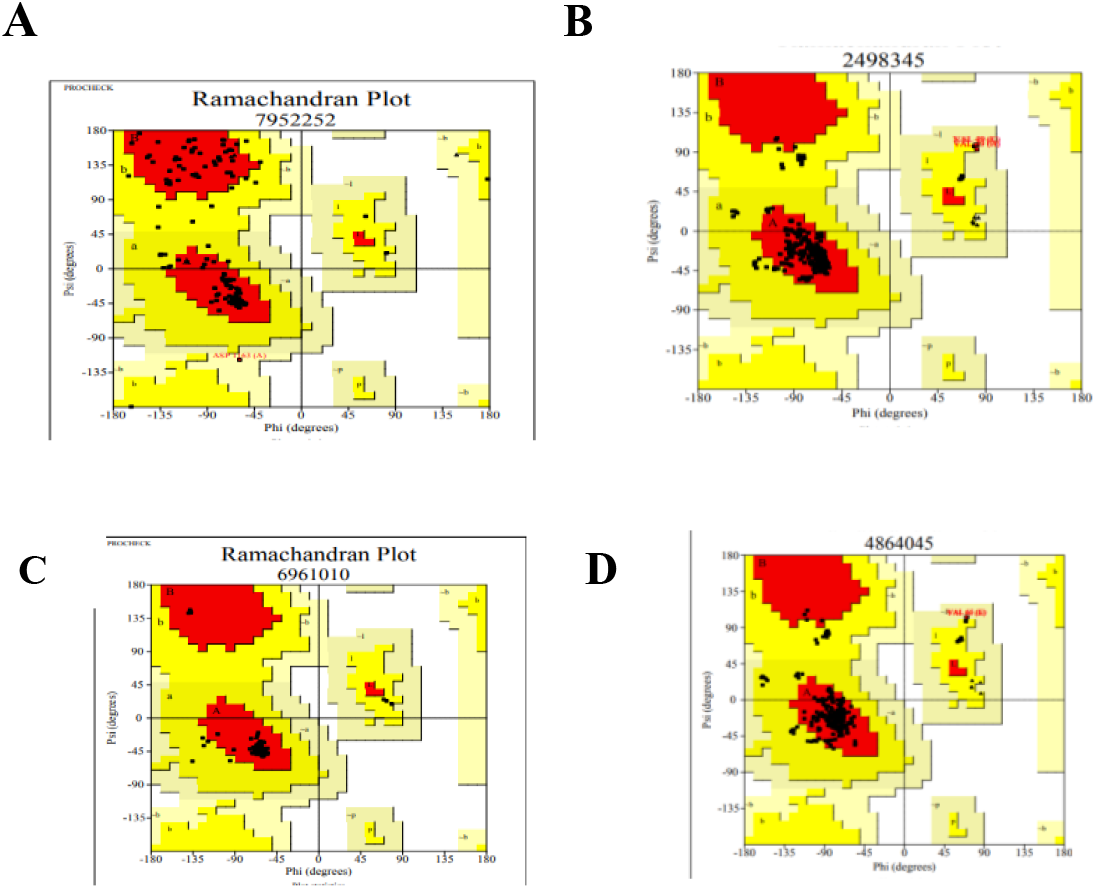
Ramachandran Plots For Structural Proteins Of Sars Cov-2 Surface Glycoprotein (A), Envelop Glycoprotein (B), Membrane Glycoprotein (C), Nucleocapsid Phosphoprotein Obtained By Procheck, Most Favored Regions Are Display In Red; Additional Allowed Regions Are Display In Yellow; Generously Allowed Regions Are Display In Beige; And Disallowed Regions Are Display In White.

The SWISS-MODEL system was used for protein structure homology modelling and alignments for all structural proteins of SARS COV 2. First, the right protein template structures in PDB were selected using the given measures: should high reporting of the template (i.e. > 60% target associated with the template) and sequence uniqueness > 35%. Instead, as an initial criterion, we used the GMQE and QMEAN4 scoring method to distinguish well from poor ones

We also carried out PROCHECK analysis to measures protein structure stereochemical consistency by comparing geometry with residue-by-residue and particularly structural geometry.

There should be more than 90 % amino acid for good quality model according to the PROCHECK standard acid resided in the most favored regions. Table 5 and Figure 4 indicate that percentage of modeled proteins residues in most favoured regions (red) is 83%—9l.4%, percentage of residues of the modeled proteins in additional allowed regions (yellow) is 8.6—15.6%, percentage of residues of the modeled proteins in generously allowed regions (beige) is 0.0—1.5%, and percentage of residues in disallowed regions (white) is 0.0%

These findings show that the models are generated overall stereochemical properties were highly stable, and that future molecular modelling studies can benefit from the models. Of those four proteins, the Ramachandran plots provide more proof of their acceptability (Figure 3).

## IV. CONCLUSIONS

The physical parameters shows that Surface protein contain highest amino acids, negatively charged residues, positively charged residues, EC and molecular weight than other proteins. The chemical factors show in proteins phenyl alanine and serine is totally absent in all structural proteins, there is a lack of histidine and glutamic acid in the protein envelope and a lack of cysteine in the nucleocapside phosphoprotein. The envelope, surface and membrane glycoprotein is rich in Leucine so have more affinity to the host cell receptor surface as stated in Luo et al, 1999. Although glycine is more in nucleocapside phosphoprotein, researchers should be able to isolate the protein with little effort along with these data and iso-electric point value. Higher molecular weight of the surface glycoprotein suggests that its tertiary structure may contain strong amino acid side chains. Instability index surface, membrane and envelop glycoprotein is stable and nucleocapsid phosphoprotein is highly unstable Leaving the surface Glycoprotein and nucleocapsid phosphoprotein all other protein are hydrophobic in nature. The prediction of trans-membrane in all structural proteins except Nucleocapsid Phosphoprotein shows the virus ability to reach the host with ease. This fairly simple method may help us understand how antivirals and vaccines could be produced against it Furthermore, as a beginning for docking studies (small and large scale) we are now offering homology models, Diverse knowledge of the molecular biology of SARS COV_2 is required to learn more.

Developing technologies will gain valuable insight into the structure of the protein in order to determine how protein disease induces, and understanding the relationship between protein-protein and protein RNA would greatly enhance our ability to develop vaccines. Meanwhile, methods of molecular simulation provide important solutions to the struggle

## Acknowledgment

All authors acknowledge and thank their respective Institutes and Universities

## AUTHORS PROFILE

**Sumaira Gulzar**

PhD scholar at Department of Biotechnology and Bioinformatics International Islamic University Islamabad Pakistan

**Saqib Husssain**

Research officer at Genome center, International Center for Chemical and Biological Sciences University of Karachi Pakistan

